# UBR-Box containing protein, UBR5, is over-expressed in human lung adenocarcinoma and is a potential therapeutic target

**DOI:** 10.1101/2020.06.03.131912

**Authors:** Kumar Saurabh, Parag P. Shah, Mark A. Doll, Leah J. Siskind, Levi J. Beverly

## Abstract

**Background:** N-end rule ubiquitination pathway is known to be disrupted in many diseases, including cancer. UBR5, an E3 ubiquitin ligase, is mutated and/or overexpressed in human lung cancer cells suggesting its pathological role in cancer.

**Methods:** We determined expression of UBR5 protein in multiple lung cancer cell lines and human patient samples. Using immunoprecipitation followed by mass spectrometry we determined the UBR5 interacting proteins. The impact of loss of UBR5 for lung adenocarcinoma cell lines was analyzed using cell viability, clonogenic assays and *in vivo* xenograft models in nude mice. Additional Western blot analysis was performed to assess the loss of UBR5 on downstream signaling. Statistical analysis was done by one-way ANOVA for *in vitro* studies and Wilcoxon paired t-test for tumor volumes *in vivo*.

**Results:** We show variability of UBR5 expression levels in lung adenocarcinoma cell lines and in primary human patient samples. To gain better insight into the role that UBR5 may play in lung cancer progression we performed unbiased interactome analyses for UBR5. Data indicate that UBR5 has a wide range of interacting protein partners that are known to be involved in critical cellular processes such as DNA damage, proliferation and cell cycle regulation. We have demonstrated that shRNA-mediated loss of UBR5 decreases cell viability and clonogenic potential of lung adenocarcinoma cell lines. In addition, we found decreased levels of activated AKT signaling after the loss of UBR5 in lung adenocarcinoma cell lines using multiple means of UBR5 knockdown/knockout. Furthermore, we demonstrated that loss of UBR5 in lung adenocarcinoma cells results in significant reduction of tumor volume in nude mice.

**Conclusions:** These findings demonstrate that deregulation of the N-end rule ubiquitination pathway plays a crucial role in the etiology of some human cancers, and blocking this pathway via UBR5-specific inhibitors, may represent a unique therapeutic target for human cancers.

## Background

Protein stability and protein turnover are key mechanisms regulating cellular processes, such as proliferation, apoptosis and senescence. The well-studied process by which cells dictate protein turnover is through the canonical ubiquitin/proteasome pathway, whereby the small protein ubiquitin is conjugated to lysine residues of substrate proteins that are to be targeted for proteasome-dependent degradation. A less studied, but related, pathway that can also regulate protein stability is through the recognition of motifs present at the N-termini of E3 ubiquitin ligase through the ‘N-end rule ubiquitination pathway’. There are seven N-recognin E3 ubiquitin ligases (UBR1-UBR7) in humans which all contain a zinc-finger domain known as a UBR-box. This domain is approximately 70 amino acids in size and functions as a recognition component in conjunction with N-degron sequences on target proteins. Through a variety of distinct mechanisms, these seven UBR-box containing proteins are involved in target recognition, ubiquitination and degradation of the proteins that have destabilized N-terminal degrons [1, 2]. Importantly, mutation and/or copy number alterations, of at least one of the seven UBR-box containing genes is found in over 25% of major cancers, including breast, bladder, cervical, lung, melanoma and serous ovarian carcinoma. Additionally, these E3 ubiquitin ligases are now known to be associated with the proteins involved in proliferation and cell cycle arrest. However, a direct link between the N-end rule ubiquitination pathway and human disease, including cancer, has yet to be demonstrated [3].

UBR5, also known as DD5, EDD, HYD, and EDD1, has been shown to be overexpressed in several solid tumors and somatically mutated in multiple cancers [3, 4, 5]. The human UBR5 gene is located at 8q22.3 downstream of the MYC locus and has 60 exons which encode an approximately 300kDa protein. There are several splice variants of UBR5 which have been reported on the NCBI database, but the clear function of these splice variants is not known [3]. UBR5 is involved in a wide array of cellular functions that include cell death, regulation of p53 and β-catenin, DNA damage response, and autophagy in multiple disease states [3, 6, 7, 8]. UBR5 was first identified in progestin-regulated genes and regulation of ERα-induced gene expression and proliferation in breast cancer cells [3, 4]. Whole genome sequencing data suggests that the 8q22 gene cluster, where UBR5 is located, is involved in cell death mediated apoptosis. In a case report of a brain metastatic sample from a pediatric lung adenocarcinoma patient, sequencing analysis reveals the presence of multiple, non-targetable mutations in several genes including the UBR5, ATM, etc [9, 10]. Thus, dysregulation in UBR5 could lead to aberration of post-transcriptional modification which could lead to the activation of multiple pathways involved in tumor progression.

Several phosphorylation sites have been reported in UBR5 and accumulating evidence suggests UBR5 might be a direct phosphorylation target of ATM-mediated DNA damage, ERK kinases and cell cycle kinases [11, 12, 13]. UBR5 has also been shown to play key roles in maintaining pluripotency of embryonic stem cells (ESC) and cellular reprograming. Further, homozygous deletion of *Ubr5* in mice results in embryonic lethality [14, 15]. Another critical cell survival and proliferation signaling pathway is through activation of AKT, which is also one of the most frequently dysregulated pathways in multiple cancers. UBR5 has been reported to interact with SOX2, a gene important in maintaining growth of ESC, as well as mediating proteolytic degradation via involvement of AKT in esophageal cancer [16]. In a recent finding, overexpression of UBR5 was shown to promote tumor growth through activation of the PI3K/AKT pathway in gall bladder cancer [5]. Although these studies all support the involvement of UBR5 in the progression of multiple cancers, the importance of this protein in lung adenocarcinoma and proliferation signaling has not been convincingly demonstrated. In this study we examine the N-end rule ubiquitination pathway, a unique biological process in lung adenocarcinoma cells, by using UBR5 as the paradigm for this complex family of proteins.

## Methods

### Cell Culture, Patient Samples and Transfection

Human embryonic kidney 293T (HEK293T) cells were procured from American Type Culture Collection (ATCC, Rockville, MD, USA) and cultured in DMEM medium (#SH30243, Hyclone, Logan, UT, USA) supplemented with 10% fetal bovine serum (#SH30070, Hyclone, Logan, UT, USA) and 1% antibiotic/antimycotic (#SV30010, Hyclone, Logan, UT, USA) at 37°C with 5% CO_2_. All lung adenocarcinoma lines were procured from ATCC and cultured in RPMI (#SH30027, Hyclone, Logan, UT, USA) supplemented with 10% FBS, 1% antibiotic/antimycotic. siRNA transfections were performed as described previously [17]. Human primary tumor and adjacent normal lung tissue samples were obtained from tissue bio-repository facility of James Graham Brown Cancer Center, at University of Louisville. Local IRB committee of the University of Louisville approved the proposed human study.

### Immunoprecipitation, Protein estimation and Western Blot

Immunoprecipitation (IP) was performed as described previously [18]. Harvested cells for each procedure (IP, transfection, infection) were lysed with 1% CHAPS lysis buffer and total protein was estimated as described previously [19]. Western blots were performed in Bolt Bis-Tris gels (#BG4120BOX, Life Technologies, Grand island, NY, USA) as per manufacturer’s protocol using antibodies from Santa Cruz, Dallas, TX, USA (GAPDH # sc47724); Bethyl, Montgomery, TX, USA (UBR5 # A300-573A, GCL1N1 # A301-843A) and Cell Signaling, Danvers, MA, USA (FLAG # 14793, DNA-PK # 4602, mTOR # 2972, RAPTOR # 2280, RICTOR # 2114, AKT # 4691, pAKT^S473^ # 4060).

### Cell viability and Clonogenic assay

Cell viability and clonogenic assays were performed as described earlier [17]. Briefly, A549 cells were cultured in 60 mm culture plates. After 24 hrs of infection with shRNA, cells were trypsinized, counted and 2000 cells were reseeded per well in 96-well plates. Cell viability was analyzed for four successive days using Alamar blue (#DAL1100 Invitrogen detection technologies, Eugene, OR. USA). At the same time following infection, 1000 cells were seeded in 6-well plates in triplicate for each condition. Cells were allowed to grow on 6-well plates for 10 days and supplemented with fresh media every two days. After 10 days, formed colonies were washed once with PBS, fixed with ethanol and stained with crystal violet for imaging and analysis.

### In Vivo Xenograft Studies

As previously described NRGS (nude mice, NOD/RAG1/2−/−IL2Rγ−/−Tg[CMV-IL3,CSF2,KITLG]1Eav/J, stock no: 024099) mice were obtained from Jackson laboratories (Bar Harbor, ME, USA) and bred and maintained under standard conditions in the University of Louisville Rodent Research Facility (Louisville, KY 40202, USA) on a 12-h light/12-h dark cycle with food and water provided ad libitum [19]. For xenograft studies, A549 cells were infected with virus particles containing shRNAs targeting UBR5 and a non-targeting (NT) control. 24hrs post-infection, cells were harvested, washed with sterile PBS and 1.25 × 10^5^ cells were suspended in 200µl PBS and delivered by subcutaneous injection in each flank of 8 male mice. For each mouse the Left flank served as the control, receiving cells treated with non-targeting shRNAs, and the Right flank received cells treated with shRNAs targeting UBR5. Four weeks post-injection mice were euthanized as per protocols, including carbon dioxide asphyxiation followed by either cervical dislocation or bilateral thoracotomy, as approved by the Institutional Animal Care and Use Committee (IACUC) of University of Louisville. The tumors were resected, and measurements made. Wilcoxon paired t-test was used to calculate the significance of difference between tumor volume.

### Reverse Phase Protein Arrays

A549 cells stably expressing Cas9 were transfected in triplicate as described above using two synthetic gRNAs of each (sgNon-targeting and sgUBR5) which were purchased Synthego (Redwood City, CA, USA). 72hrs post-transfection, the cells were harvested, washed with PBS then frozen until further processing. Frozen cell pellets were then shipped to MD Anderson Cancer Center, Houston, TX, USA for protein isolation and Reverse Phase Protein Arrays (RPPA) analysis.

### Statistical Analysis

All statistics were performed using GraphPad Prism 8 software. Unless otherwise specified, significance was determined by one-way ANOVA, using a cut off of p < 0.05.

## Results

### UBR5 is altered in human lung adenocarcinoma

Previous reports and database searches identified UBR5 as a gene that is mutated, amplified and over-expressed in many human cancers. Recent data on lung adenocarcinoma from The Cancer Genome Atlas (TCGA) reveals that UBR5, a UBR-box containing 2799 amino acid protein, is either altered by mutation, increased copy number or amplified in more than 20% of the samples (Fig. 1a & 1b). To initially establish the UBR5 expression levels in cancer cell lines and primary human patient samples, we examined the basal protein level of UBR5 in various lung adenocarcinoma cell lines being cultured in our laboratory (Fig. 1c). A wide variability in the UBR5 protein level was detected in these different cell lines by Western blot analysis, but interestingly, UBR5 was not detectable in IMR90 cells (a non-transformed lung fibroblast cell line). In addition, we performed Western blot studies on 15 primary resected non-small cell lung cancer samples (T) and their adjacent normal lung tissue (N). When compared to the adjacent normal lung, UBR5 expression levels were uniformly higher in the cancer samples. (Fig. 1d).

**Figure 1:**
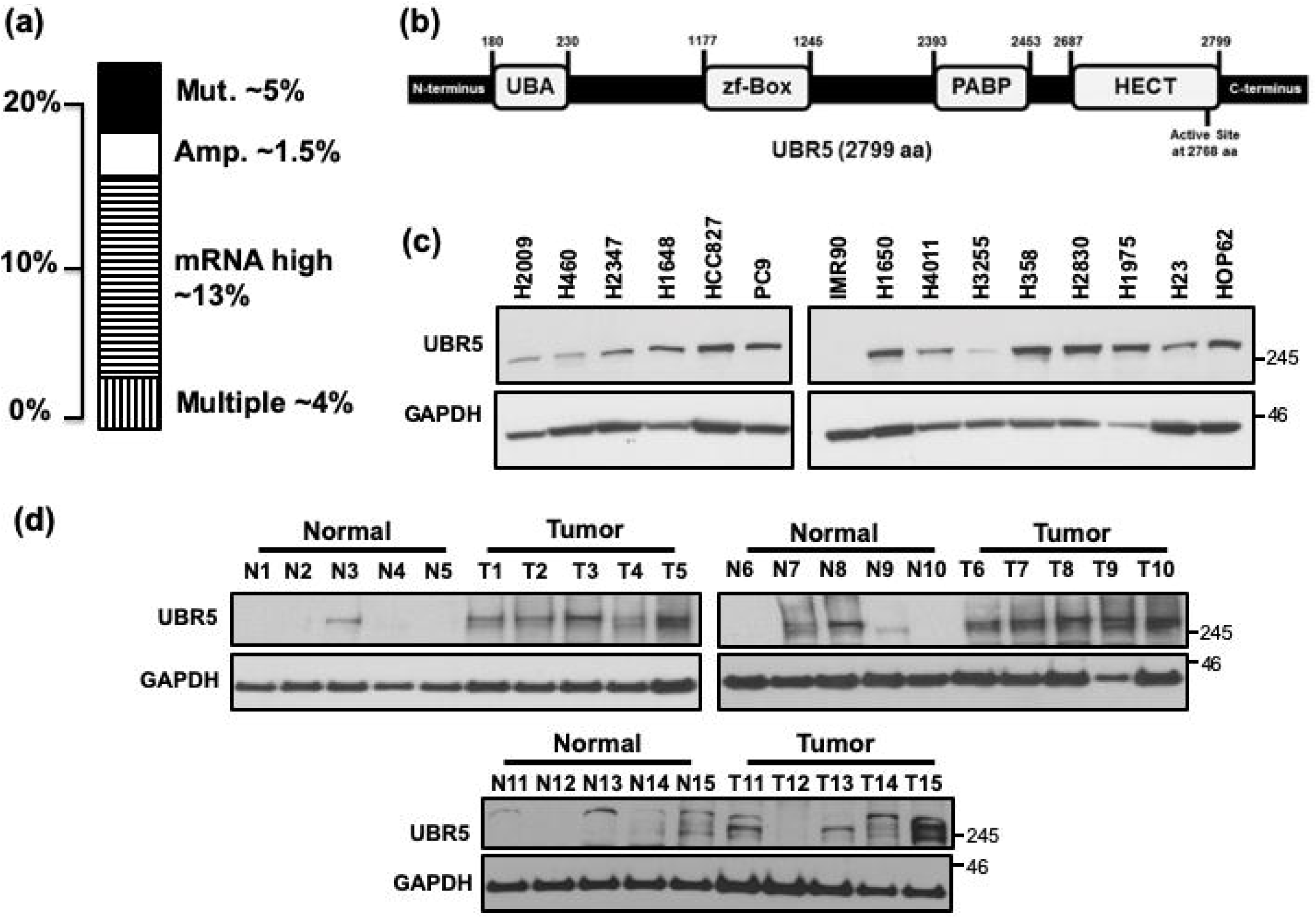
UBR5 is altered in human lung adenocarcinoma. (a) UBR5 is altered by mutation or increased copy number in 10% of human lung adenocarcinoma analyzed by TCGA. (b) Schematic of UBR5 protein. UBR5 is a 2799 amino acid protein with an N-terminal UBA (ubiquitin-associated) domain, a middle zf-Box (zinc finger UBR-box) domain and two C-terminal domains, PABP (poly[A]-binding protein) & HECT (homology to E6-AP carboxyl terminus). (c) A wide variability in the UBR5 protein level was observed in various lung adenocarcinoma cell lines; Tubulin was used as a loading control. (d) The level of UBR5 protein was found to be highly elevated in nearly all primary human lung tumor samples (T) compared to normal adjacent lung (N), N1 represents the normal adjacent to T1, N2 is the normal adjacent to T2, etc.; GAPDH was used as a house keeping gene. Western blot images were cropped from original scans, which are provided in additional materials.

### UBR5 interacts with multiple proteins

To gain insight into the possible mechanism(s) by which UBR5 regulates cellular signaling pathways involved in apoptosis and cell survival, we performed immunoprecipitation of UBR5, followed by mass spectrometry (IP/MS) analysis to identify UBR5 interacting proteins (Fig. 2a). We sorted the MS data based on highest unique peptide count which revealed several interacting proteins involved in multiple cellular pathways like biosynthesis of amino acids, inflammation, differentiation, DNA replication, apoptosis, etc. Some of the proteins which caught our attention were GCN1L1 (a positive activator of the EIF2AK4/GCN2 protein kinase activity in response to amino acid starvation); CASP14 (a non-apoptotic caspase involved in epidermal differentiation); ANXA1 (involved in the innate immune response as an effector of glucocorticoid-mediated responses and regulator of the inflammatory process); and SLC25A5/6 (catalyzes the exchange of cytoplasmic ADP with mitochondrial ATP across the mitochondrial inner membrane) (Fig. 2a). To further validate this MS data, we transiently transfected HEK293T cells with a UBR5-Flag construct, then pulled down UBR5-interacting proteins using anti-Flag beads (Fig. 2b). Interestingly, Western blot analysis of these Flag-IP samples revealed that UBR5 was interacting with proteins of the mTOR complex including Raptor/Rictor. Additionally, UBR5 co-immunoprecipitated with DNA-PK (DNA damage pathways), GCN1 (a translational activator), CDK1 (cell cycle regulator), and AKT (known roles in cell proliferation and migration) (Fig. 2a).

**Figure 2:**
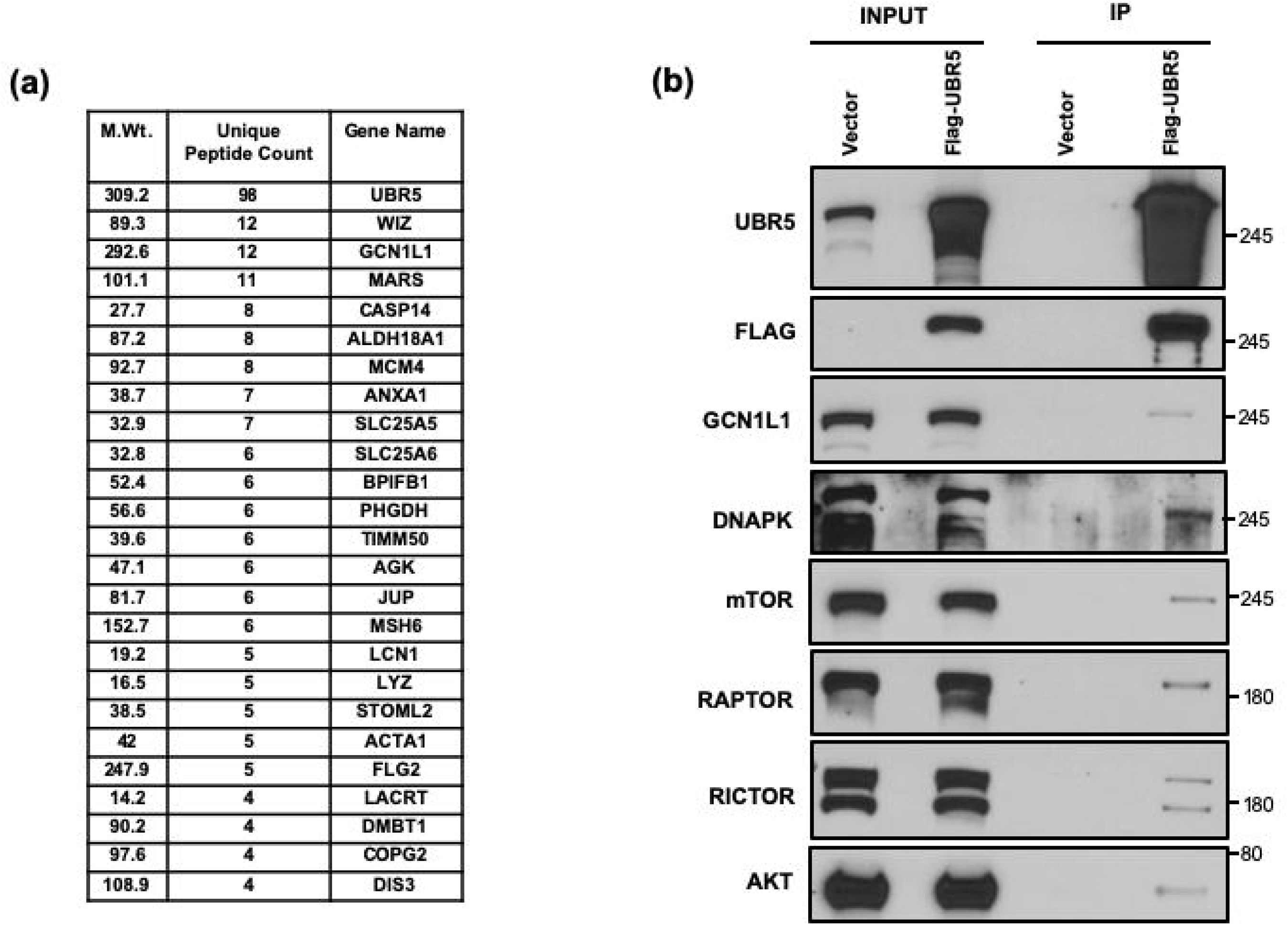
UBR5 interacts with multiple proteins. (a) An abbreviated list of proteins that were identified as UBR5 interacting proteins. Unique peptide count; number of distinct peptides identified by MS/MS in UBR5 immunoprecipitates (IP). Many secretory and anti-inflammatory proteins are found to be interacting with UBR5. (b) HEK293T cells were transiently transfected with FLAG-tagged UBR5 followed by IP by anti-FLAG antibody and Western Blot analysis. UBR5 interact with proteins involve in DNA damage and mTOR pathway. Western blot images were cropped from original scans, which are provided in additional materials.

### UBR5 interacts with total and phosphorylated AKT

The interaction with mTOR components and AKT warranted further investigation since AKT is involved in various cellular processes and is known to promote survival and growth. AKT is also a key component of mTOR signaling transduction which has been shown to crosstalk with multiple signaling pathways. Strikingly, Western blot analysis of IP samples confirmed that phosphorylated and total AKT both interact with UBR5 (Fig. 3a). Given that UBR5 is highly expressed and/or mutated in lung cancer, we were interested in determining the outcome of UBR5 loss and AKT status. To this end we utilized multiple techniques to reduce the protein levels of UBR5 in lung adenocarcinoma cells. A549 cells were infected with lentivirus containing multiple shRNA molecules designed to target different coding regions of UBR5. Loss of UBR5 resulted in a robust decrease of phosphorylated AKT but shows no change in expression of total AKT (Fig. 3b). To further explore these findings, we generated cell lines of A549 and H460 cells that stably expressed Cas9, then transiently transfected these cells with two synthetic gRNAs designed to target UBR5 or the corresponding non-targeting control gRNAs. Interestingly, both cell lines showed similar decreases in phosphorylated AKT when the levels of UBR5 were reduced (Fig. 3c). This reduction in phosphorylated AKT was consistent for all methods and in multiple cell lines, thus UBR5 is likely a regulator of AKT phosphorylation status.

**Figure 3:**
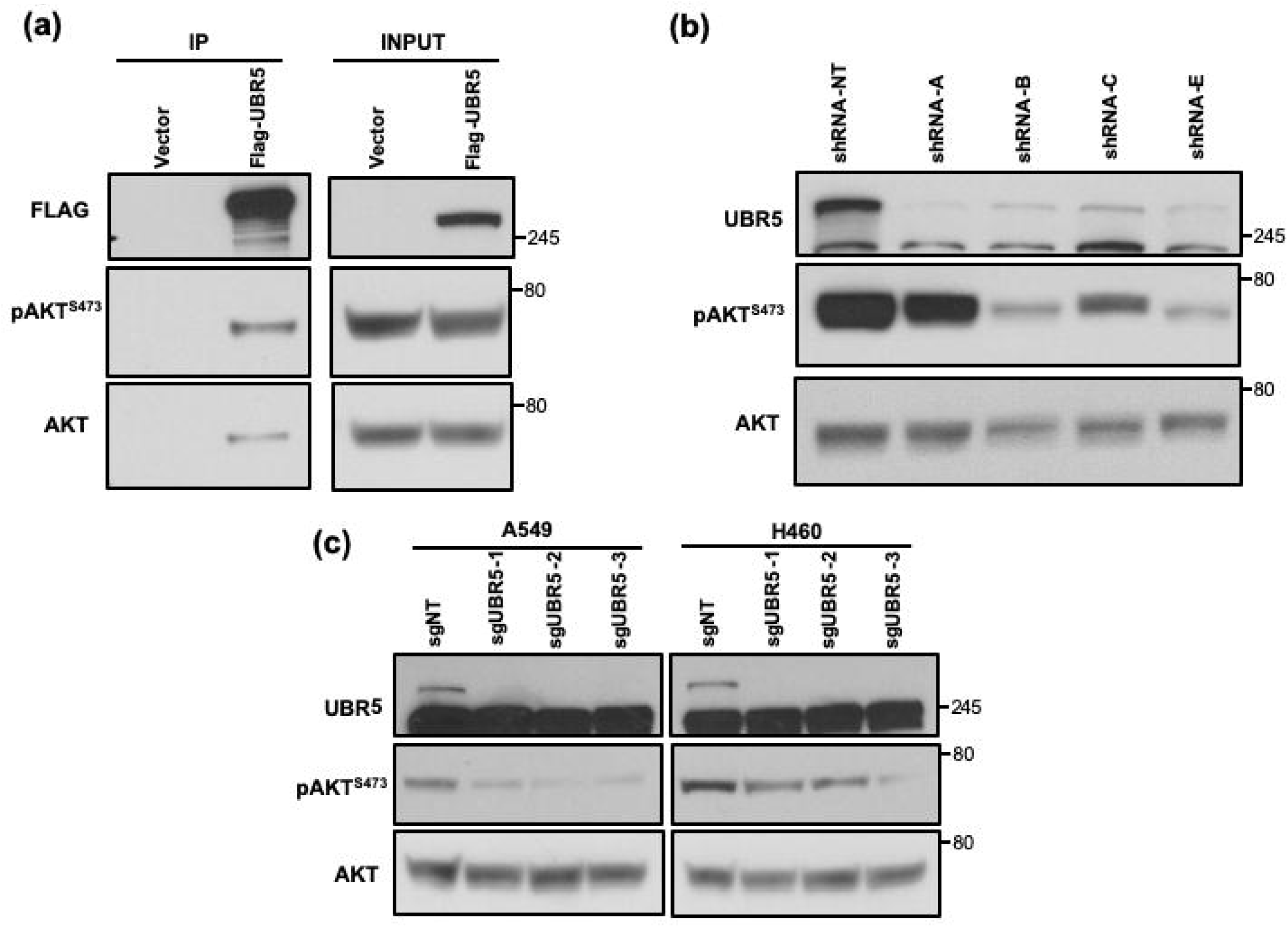
UBR5 interacts with AKT and regulate its activity. (a) HEK293T cells were transiently transfected with FLAG-tagged UBR5 followed by IP by anti-FLAG antibody and Western Blot analysis. (b) A549 cells were infected with the lenti-viruses from multiple shRNA construct against UBR5. (c) Lung adenocarcinoma cell lines which are stably expressing Cas9 were transiently transfected with synthetic gRNAs against UBR5. Western blot images were cropped from original scans, which are provided in additional materials.

### Loss of UBR5 has global impacts on multiple cellular pathways

The UBR5-AKT interaction prompted us to ask which other signaling pathways might be impacted by the loss of UBR5. To address this question, we transiently transfected triplicate cultures of a Cas9 expressing A549 cell line with gRNAs designed to target UBR5 or the corresponding non-targeting control gRNAs. Cells were harvested and frozen then samples were sent in triplicate for RPPA analysis at MD Anderson Cancer Center, Houston, TX, USA (Fig. S1). This technique provides a high throughput approach to determining protein expression levels under various conditions. To calculate the change in protein expression as a result of UBR5 loss, protein levels from cells treated with non-targeting gRNAs were set as the baseline value of one. This was compared to the average of six separate readings from cells treated with two distinct gRNAs targeting UBR5 and the fold change for each protein was calculated. We then filtered the data to create lists consisting of the top 20 downregulated and top 20 upregulated proteins resulting from the loss of UBR5. This in-depth analysis allowed us to identify multiple cellular pathways that were directly impacted by the loss of UBR5. In the absence of UBR5, many of the downregulated proteins are known to be involved in processes like endocytosis, extracellular matrix organization, cell migration and cell survival (Fig. S1). On the other hand, proteins like p21, PAR1, NQO1, S6 kinase and CD31, were among the top 20 upregulated proteins in the absence of UBR5. These proteins are known to be highly regulated during processes like apoptosis, DNA damage and cell cycle arrest (Fig. S1).

### UBR5 deficient A549 cells show decreased cell growth and clonogenic potential

Since UBR5 is highly expressed and/or mutated in lung cancer, our cancer cell lines offered a viable model for focusing on the outcome of UBR5 loss. Interestingly, multiple shRNAs targeting UBR5, resulted in decreased cell growth in lung adenocarcinoma cell lines (Fig. 4a). The knockdown of UBR5 resulted in a robust effect with approximately 90% loss of cell viability by the end of day four (Fig. 4a). We extended this further to determine the effect of UBR5 loss on colony formation. To this end, A549 cells were infected with multiple shRNAs targeting the coding region of UBR5, then seeded at 1000 cells per well for 14 days (Fig. 4b). This clonogenic assay approach shows there were significantly reduced colonies formed from the cells deficient for UBR5 proteins as compared to the cells transduced with non-targeting shRNAs (Fig. 4b & 4c).

**Figure 4.**
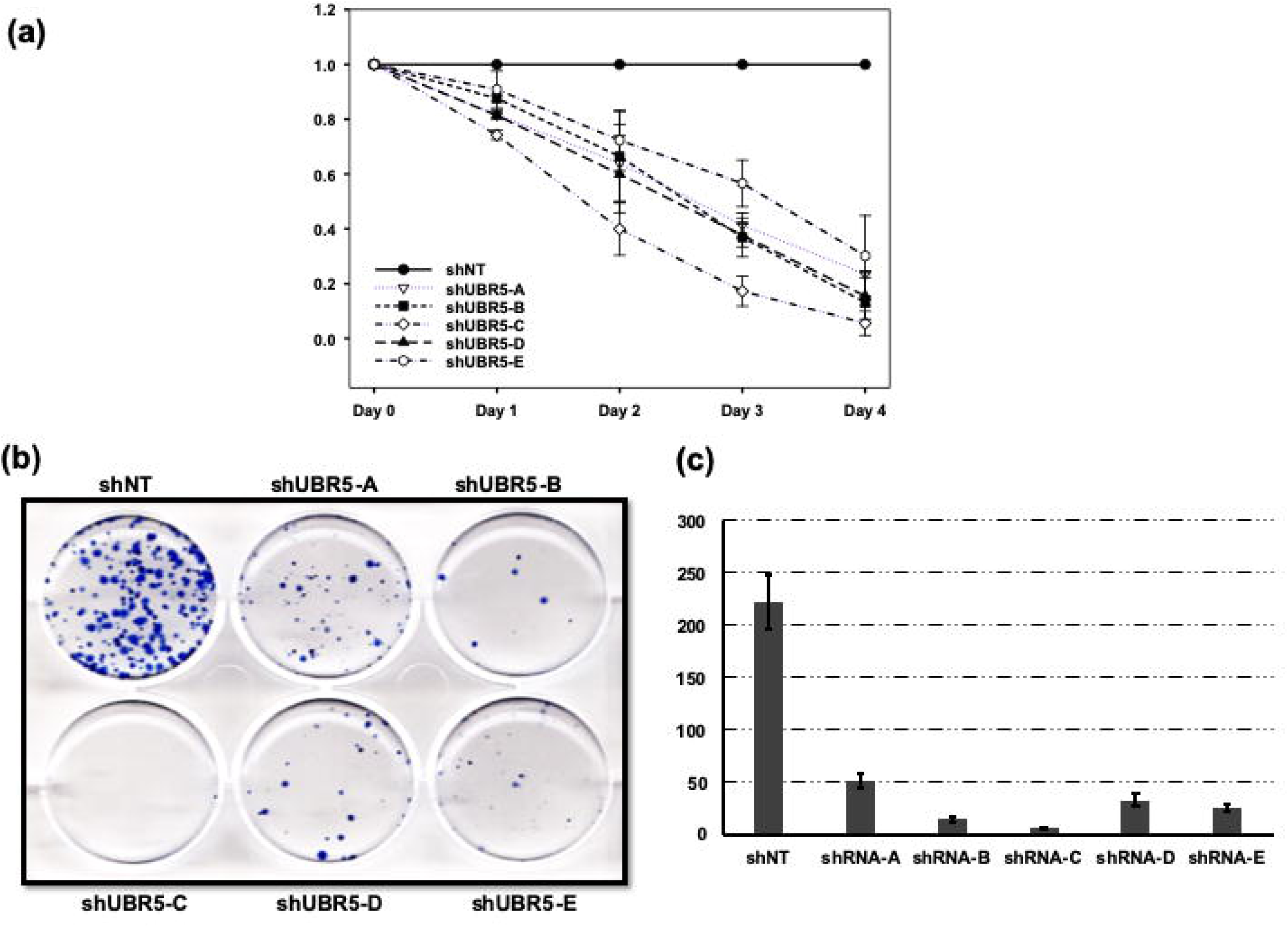
UBR5 deficient A549 cells show decreased cell viability and clonogenic potential. (a) A549 cells were infected with the lenti-viruses from multiple shRNA construct against UBR5 and were cultured for 4 days. Alamar Blue readings were recorded every 24 hours and relative cell viability of UBR5 deficient cells were compared to control cells on each day. (b) A549 cells were infected with shRNA against UBR5 and 1000 cells were cultured in 6-well plate for 10 days. Colonies were fixed in methanol and stained with crystal violet. (c) Quantitative evaluation of clonogenic assay. Representative bar graph showing number of colonies formed per 1000 cells

### Loss of UBR5 results in reduction of tumor volume in NRGs mice

To further understand the possible clinical significance of the loss of UBR5 in cancer cells, we directed our studies to explore *in vivo* tumor formation in nude mice. A549 cells were infected with lentiviral constructs that either targeted UBR5, or the corresponding non-targeting controls. After harvesting, these cells were subcutaneously injected in the flanks of NRGs (immune deficient) mice. Cells treated with non-targeting shRNAs were injected in the Left flank (Control group) and cells treated with UBR5-specific shRNAs were injected in the Right flank (Fig. 5a). Consistently, tumors that developed from the non-targeting shRNA treated cells were larger in size and weight when compared to the tumors that developed from the UBR5-specific shRNAs (Fig. 5a). We further quantified the volume of each tumor as shown (Fig. 5a, 5b). The data indicate that tumors generated from cells treated with shRNAs targeting UBR5 are significantly smaller in volume, irrespective of the coding region being targeted (Fig. 5b). We further confirmed the significance of this tumor volume difference by performing a Wilcoxon paired t-test (Fig. 5c). These results strongly suggest that the loss of UBR5 is critical to tumor progression and could be clinically relevant to lung cancer studies.

**Figure 5.**
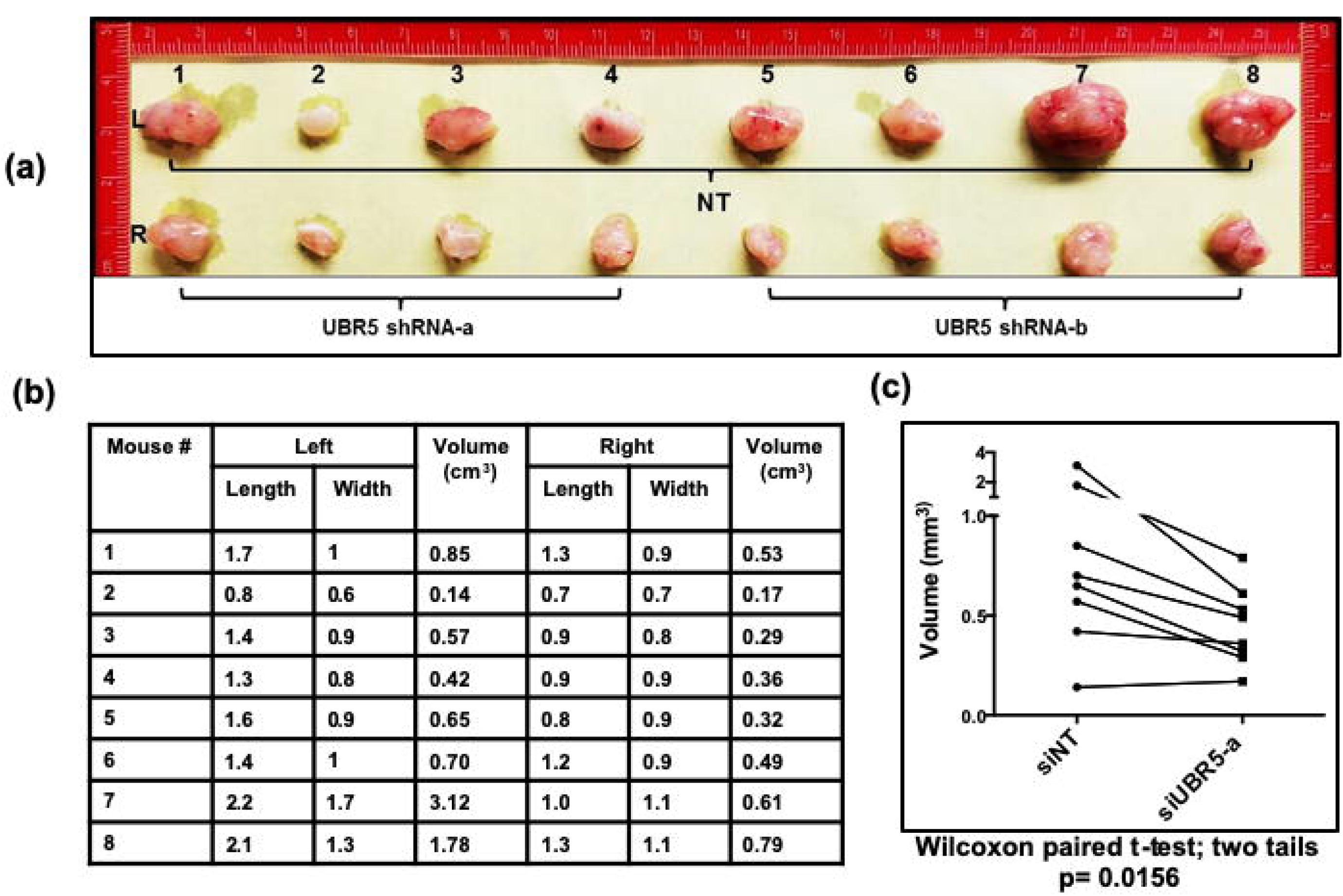
Loss of UBR5 results in reduction of tumor volume in NRGs mice. (a) A549 cells were infected with the lenti-viruses from multiple shRNA construct against UBR5. 24hrs post infection, cells were harvested and 1.25 × 10^5^ cells were subcutaneously injected in mice flanks (L-NT shRNA & R-shRNA against UBR5). (b) Mice were euthanized 4 weeks after injection and tumor were measure. (c) Wilcoxon paired t-test were used to calculate the significance of difference between tumor volume.

## Discussion

The results reported here allowed us to advance our understanding of how UBR5, as part of the N-end rule ubiquitination pathway, works to regulate basic biological processes, and what involvement it might have in cancer initiation, progression and maintenance. Importantly, cancers with mutations and/or amplification of genes involved in N-end rule ubiquitination, may represent a unique subset of cancers that are exquisitely susceptible to novel therapeutic interventions [20, 21, 22, 23]. Our data show that UBR5 is highly expressed in multiple lung adenocarcinoma cell lines but is present at reduced levels or absent in normal cell lines. Western blot analysis data further confirms high expression levels of UBR5 protein in primary resected non-small cell lung cancer samples when compared to the adjacent normal lung tissue from the same patient. Previous reports and database searches identified UBR5 as a gene that is mutated, amplified and over-expressed in many human cancers [24, 25]. Since human UBR5 was initially identified to be somatically mutated in breast cancer cell lines, several studies have subsequently shown UBR5 to be dysregulated in a wide array of other cancers. A recent study shows that UBR5 was responsible for polyubiquitination of anterior gradient 2 (AGR2), a protein that was first identified as being downregulated in breast cancer cell lines via sequencing data [4]. In support of this, proteasome inhibition also results in suppression of AGR2 transcription by downregulation of E2F1 [4, 26]. The TCGA data from lung adenocarcinoma reported in this study, suggests that UBR5 is either altered, or the gene copy number is significantly increased, in several patient samples. These amplifications are most likely in the form of allelic imbalance resulting in increased mRNA copy number of UBR5 [25]. Our findings further support previous pathological studies looking at the gene cluster on chromosome 8q22 (where UBR5 is located) showing it is amplified/mutated in lung cancer [9, 10]. The fusion of locus 8q22 and zinc finger protein 423 (ZNF423) on 16q12 was also identified in head and neck cancer primary tumors where downregulation of this fusion inhibits cell proliferation in nasopharyngeal carcinoma [27]. When considering the multiple mutations reported to date for UBR5, many of them are found to be associated with the conserved C-terminal cysteine of the HECT domain. This domain serves as a recognition site for the target protein and alteration of this domain is associated with disrupted UBR5 ligase activity [28, 29].

Over-expression of UBR5 contributes to the genesis of human cancer. Our IP/MS data further reveals UBR5 interacting with multiple proteins known to be involved in metabolism, inflammation, DNA replication, differentiation and apoptosis. For example, our data demonstrates that Annexin A1 (ANXA1) interacts with UBR5. ANXA1 is a direct regulator of NFkB signaling via binding to p65 and regulating p65-induced transcription. Furthermore, increased expression of ANXA1 has been shown to induce apoptosis in some cell types [30, 31]. Additional examples of UBR5 interacting proteins from our MS data include translational activators (GCN1L1), non-apoptotic caspase (CASP14) and mitochondrial ATP transporters (SLC25A5 and SLC25A6). The SLC25 family of proteins has been shown to be overexpressed in various cancers and involved in ADP/ATP exchange between mitochondrial matrix and cytosol [32]. UBR5 interaction with SLC25 family member proteins further suggests that disruption in UBR5 expression could result in mitochondrial dysfunction and prevention of apoptosis in disease state. Previous studies have reported that UBR5 interacts with alpha4, a component of the mTOR pathway, in human MCF-7 breast cancer cell line [33]. The mTOR/AKT pathway/signaling cascade is crucial in the regulation of cellular proliferation, inhibition of apoptosis and metabolism. Proteins in this pathway are often reported to be mutated or amplified in lung adenocarcinomas. Also, the mTOR complex is known to activate AKT through phosphorylation at Serine 473 [34, 35]. Our IP data suggests a direct interaction of UBR5 with proteins of the mTOR complex, including Raptor and Rictor. Further experiments would be needed to elucidate the nature of these interactions. We have also observed that UBR5 interacts with DNAPK, a molecule known to be activated during DNA damage responses, which further supports a role for UBR5 in ATM and DNA-PK mediated H2AX-phosphorylation, as well as cancer development and progression [11, 36].

The activation of the AKT pathway is known to play a role in cell survival and cell proliferation in various types of cancers. In a similar study, UBR5 was shown to be overexpressed in gall bladder cancer and downregulation of UBR5 inhibited the cell proliferation of relevant cancer cell lines [5]. The data reported by Zhang, et al., suggests that loss of UBR5 results in increase of PTEN, a tumor suppressor gene which acts as a negative regulator of the PI3K/AKT pathway in gall bladder cancer [5]. As reported here, we also observed a robust decrease in phosphorylated AKT at Serine 473 using multiple methods of reducing UBR5 levels in lung adenocarcinoma cell lines. Additionally, UBR5 was previously shown to interact with SOX2 and mediated its proteolytic degradation in ESC. SOX2 was shown to be overexpressed in esophageal cancer and inhibiting AKT stabilizes SOX2 expression in esophageal squamous cell carcinomas [16]. UBR5-mediated ubiquitination of citrate synthase has also been shown to play a role in AKT activation during hypoxic conditions. Loss of citrate synthase via ubiquitination by UBR5, leads to an accumulation of citrate in the cytosol during hypoxia, which leads to the activation of AKT signaling resulting in increased invasion and metastasis of breast cancer cells [37]. Excitingly, as reported here, we have found that UBR5 interacts with total and phosphorylated AKT, and the loss of UBR5 results in decreased phosphorylation of AKT. These findings suggest that post-transcriptional modification of proteins by UBR5 could be responsible for normal signaling and disruption of this UBR5-mediated regulation could lead to increases in the activation of AKT signaling, further enhancing the invasiveness and metastatic properties of cancer cells.

UBR5 has been shown to promote cell proliferation and inhibit apoptosis. Our data clearly demonstrate that loss of UBR5 leads to rapid and robust loss of cell viability and clonogenic potential in lung adenocarcinoma lines. In agreement with our results, a study by Ji, et al., showed that the loss of UBR5 in colon cancer cells was shown to decrease cell proliferation and could involve the degradation of p21, a cyclin-dependent kinase inhibitor, by polyubiquitination [38]. Similar regulations by UBR5 have been shown in gastric and colorectal cancer where increased ubiquitination by UBR5 destabilizes tumor suppressor genes leading to a reduction in the stability of these proteins [39, 40]. Collectively, these finding suggest that UBR5 disruption is involved in the genesis of human cancer through its ability to regulate the stability of multiple proteins involved in inhibiting apoptosis. As reported here, our RPPA data highlights a variety of proteins whose expression levels were altered by the loss of UBR5. This analysis reveals the critical role that UBR5 might play in maintaining extracellular matrix organization, promoting invasion and migration, and increasing cell survival in lung adenocarcinoma cells. Loss of UBR5 could lead to the destabilization of proteins which are responsible for these critical cell survival and proliferation processes. Additionally, proteins like p21 and phosphorylated S6, were found to be upregulated after loss of UBR5, which further supports a critical role for UBR5 [35, 38]. Further validation of the many proteins revealed in this study to be impacted by the loss of UBR5, has the potential to uncover novel targets for each of the signaling pathways discussed. Loss of UBR5 has been shown to be embryonic lethal. A knock-out mouse of UBR5 was reported previously, but the mice did not survive past embryonic day 10.5 because of failed yolk sac development [14]. As multiple human cancers exhibit increased expression levels of UBR5, we were prompted to investigate what impact the loss of UBR5 might have on cancer cells that have become reliant on this protein for survival. For these experiments, we used an established xenograft immunodeficient mouse model (NRGs). Remarkably, cells lacking UBR5 resulted in a significant reduction of tumor volumes as reported here. This finding further supports previous results reported in xenograft mouse models in which subcutaneously injected breast and gallbladder cells lacking UBR5 yielded smaller tumors [4, 5]. Collectively, these data suggest that UBR5 plays an essential role in cell survival and the loss of UBR5 has a tumor suppressive role in multiple human cancers. Therefore, lung cancer patients might not be the only subset of cancer patients who would benefit from inhibition of the N-end rule ubiquitination pathway.

## Conclusions

This study demonstrates that continued expression of UBR5 is required for the survival and proliferation of human lung cancer but not normal lung fibroblasts. UBR5, a key molecule in N-end ubiquitination pathways, was shown to interact with a wide range of proteins involved in cell proliferation, DNA damage, differentiation and apoptosis. AKT phosphorylation is known to play a vital role in various cellular processes which promote survival and growth in response to extracellular signals. AKT regulation is a key component of mTOR signaling transduction which has recently been shown to intersect with other signaling pathways. We confirmed that AKT interacts with UBR5 and this interaction could result in loss of AKT phosphorylation which might further impact mTOR-mediated signaling pathways to inhibit apoptosis.

Currently there are still gaps in our knowledge of the role(s) UBR5 has in normal cell biology, as well as how mutations in this gene might impact tumorigenesis. Elucidating these aspects of UBR5 biology, and the N-end rule pathway overall, could reveal novel interventions for inhibiting the progression of multiple types of human cancers. Identifying patients with cancers exhibiting increased expression levels of the genes involved in N-end rule ubitquitination, like UBR5, might afford these patients access to cytotoxic therapeutics that better target their subset of cancer. We conclude that deregulation of the N-end rule ubiquitination pathway plays a causal role in the etiology of some human cancers and blocking this pathway, via UBR5-specific inhibitors, is a unique therapeutic target for the eradication of human cancers.

## List of Abbreviations

AKT: Protein kinase B (PKB);
ERα: Estrogen receptor alpha;
ERK: Extracellular signal-regulated kinases;
SOX2: SRY(sex determining region Y)-box 2;
PI3K: Phosphoinositide 3-kinases;
EIF2AK4: Eukaryotic translation initiation factor 2-alpha kinase 4;
CASP14: Caspase 14;
ANXA1: Annexin A1;
mTOR: mammalian target of rapamycin;
DNA-PK: DNA-dependent protein kinase;
CDK1: Cyclin-dependent kinase 1;
Cas9: CRISPR associated protein 9;
p21: CDK inhibitor 1;
PAR1: Prader-Willi/Angelman region-1;
NQO1: NAD(P)H dehydrogenase (quinone) 1;
CD31: Platelet endothelial cell adhesion molecule (PECAM-1);
E2F1: Transcription factor E2F1;
HECT: homology to E6-AP carboxyl terminus;
H2AX: H2A histone family member X.

## Declarations

### Ethics approval and consent to participate

Human primary tumor and adjacent normal lung tissue samples were obtained from the tissue bio-repository facility of James Graham Brown Cancer Center, at University of Louisville, Louisville, Kentucky. The IRB committee of the University of Louisville approved the proposed human study. All animal experiments have been approved with the Institutional Animal Care and Use Committee (IACUC # 11090) of University of Louisville.

### Consent for publication

Not applicable

### Availability of data and materials

The datasets used and/or analyzed during the current study are available from the corresponding author upon reasonable request.

### Authors’ contributions

KS and PPS did the experiments. MAD generated clones of Cas9 expressing lung adenocarcinoma cell line. KS, LJS and LJB conceived studies, did the analysis and wrote the manuscript. All authors have read and approved the manuscript.

## Acknowledgements

We are grateful to the Beverly and Siskind lab members for continuous support and sharing resources. We thank RPPA Core Facility at MD Anderson Cancer Center, University of Texas, Houston, TX, USA for providing RPPA data and analysis.

## Figure Legends

**Figure S1.**
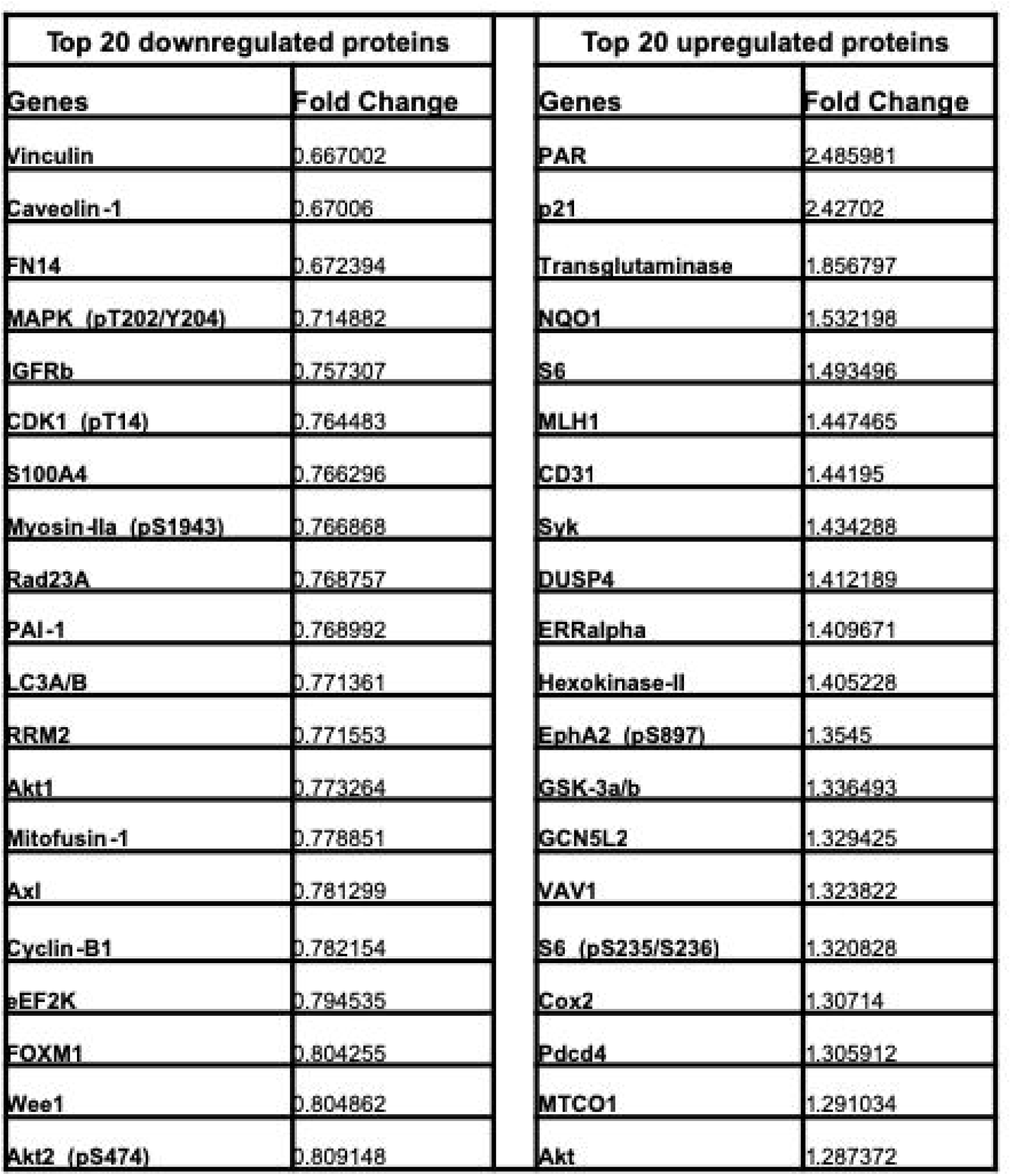
Cas9 expressing A549 cells were transiently transfected with two gRNA against UBR5. Samples were prepared in triplicate and send for RPPA analysis at MD Anderson Cancer Center, Houston, TX, USA. Each reading is average of 6 numbers, where gRNA targeting NT were considered as 1 and each column shows relative fold change of protein level.

